# The sex determination gene *doublesex* is essential for female-specific flight and hearing properties in *Aedes aegypti* mosquitoes

**DOI:** 10.64898/2026.08.28.747963

**Authors:** YuMin M Loh, WenWei Loh, Jia Xin Yap, Yuichiro Tsuchiya, Tai-Ting Lee, Yifeng YJ Xu, Daniel F Eberl, Ming Li, Omar S Akbari, Matthew P Su, Azusa Kamikouchi

**Affiliations:** Graduate School of Science, Nagoya University, Nagoya 464-8602, Japan; Institute of Transformative Bio-Molecules (WPI-ITbM), Nagoya University, Nagoya 464-8601, Japan; Department of Biology, University of Iowa, Iowa City, IA, USA; School of Biological Sciences, Department of Cell and Developmental Biology, University of San Diego, La Jolla, California, USA; Department of Parasitology, Zhongshan School of Medicine, Sun Yat-sen University, Guangzhou, 510080, China; Institute for Advanced Research, Nagoya University, Nagoya 464-8601, Japan

## Abstract

The acoustic communication system supporting reproduction in disease-transmitting mosquitoes is sexually dimorphic, from the sounds that mosquitoes make to the properties of their flagellar ears. The molecular mechanisms shaping these dimorphisms, however, remain unclear. Here, we explored the contributions of the female-specific isoform of the sex determination gene *doublesex* (*dsxF*) in specifying female-unique properties in the acoustic communication system of *Aedes aegypti* mosquitoes. CRISPR-Cas9 based generation of a novel *dsxF* knockout line found that *dsxF* mutant females were flightless due to lack of expression of a female-specific myosin gene, *myo-fem*, in the thorax. We found that DsxF directly binds to four different enhancer sites of *myo-fem* to promote gene expression. We observed that *dsxF* female ears exhibited intersex neuroanatomical characteristics and contained a more extensive auditory efferent network than control females. At the peripheral hearing function level, *dsxF* female ears displayed intersex tuning frequencies associated with increased expression of hearing genes in *dsxF* female ears compared to control females. We identified three hearing genes transcriptionally inhibited by DsxF *via* direct binding. DsxF is a bifunctional transcription factor, activating female-specific gene expression in thoraces whilst repressing male-biased gene expression in ears to coordinate female-unique features across sensory and motor systems.

## Introduction

In disease-transmitting mosquitoes, sexually dimorphic Wing Beat Frequencies (WBFs) facilitate male attraction to female WBFs (“phonotaxis”) ^1^. In *Aedes aegypti* (*Ae. aegypti*) mosquitoes, male WBFs are about 300Hz higher than female WBFs (Fig 1A) ^2^, potentially linked to sex-biased expression of flight-related genes in the thorax.

**Figure 1:**
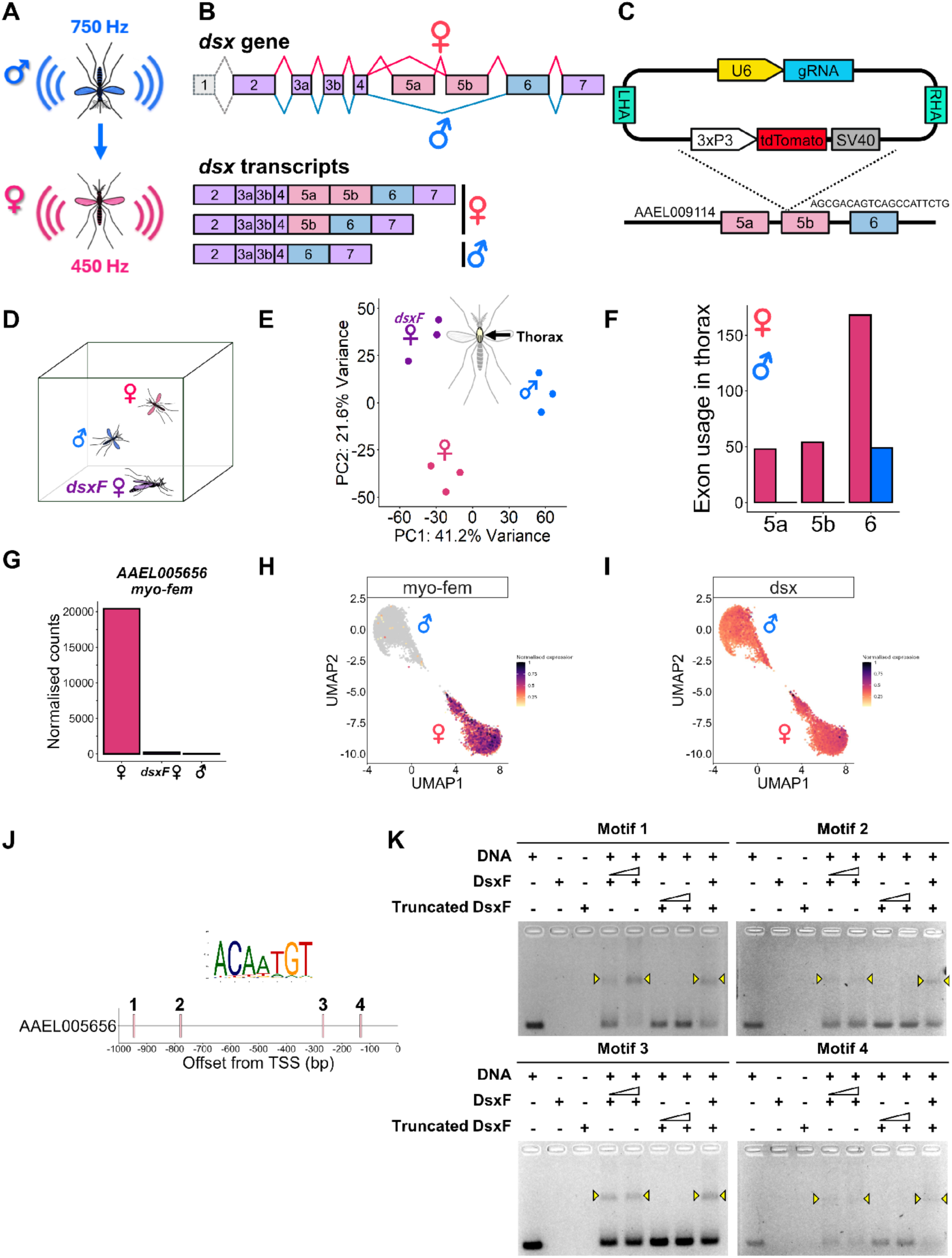
DsxF activates *myo-fem* expression *via* direct binding to support female flight (A) Male attraction to female flight sounds relies on sexually dimorphic Wing Beat Frequencies (WBFs). (B) Sex-specific alternative splicing of *doublesex* (AAEL009114) in *Aedes aegypti*, with exons 5a and 5b being specific to female *dsx* transcripts only. (C) Schematic of *dsxF* mutant generation using CRISPR-Cas9. gRNA was designed to target a site within exon 5b of *doublesex* gene, and homology-directed repair was used to insert a 3xP3-tdTomato-SV40 fragment into the target site. gRNA = gRNA scaffold, LHA = Left Homology Arm, RHA = Right Homology Arm, SV40 = SV40 transcriptional terminator, U6 = *Ae. aegypti* U6 promoter. (D) Schematic of *dsxF* mutant female flightless phenotype. (E) PCA of thorax bulk RNA-sequencing data for control females, *dsxF* mutant females and control males. (F) *dsx* splicing in control female and control male thoraces analysed based on thorax bulk RNA-sequencing data. Red and blue represent female and male exon usage respectively. (G) Expression level of *myo-fem* (AAEL005656) in control female, *dsxF* mutant female and control male thoraces in thorax bulk RNA-sequencing dataset. (H) Expression pattern of *myo-fem* in published thorax snRNA-seq dataset ^17^. See Fig S2C for sex-specific muscle cells, identified based on the sex of the thorax tissue samples. Color intensity represents normalized UMI-count. (I) Expression pattern of *dsx* in published thorax snRNA-seq dataset ^17^. (J) Occurrence of Dsx-binding motif in the promoter region (1kbp upstream of translational start site) of *myo-fem*. (K) EMSA data showing protein-DNA interactions between full-length DsxF protein or truncated DsxF protein without DM domain and the four 100 bp promoter fragments of *myo-fem* that each bears a centered Dsx-binding motif. Interactions represented as band shifts were observed only between full-length DsxF protein and target DNA sequences. See Fig S2D for whole gel images.

Beyond the sound source (WBFs), the sound receivers (flagellar ears) of mosquitoes are also highly sexually dimorphic ^3^. Mosquito flagellar ears consist of a hairy flagellum coupled to its base the Johnston’s organ (JO), the site of auditory mechanotransduction ^1,4^. Male flagella are more plumose than female flagella and male JOs also house two-fold more mechanosensory neurons than female JOs ^4^.

Functionally, male and female ears have distinct tuning frequencies tailored to extract different acoustic information from their near-field surroundings. Sexually dimorphic tuning frequencies support sex-specific hearing needs, such as the tuning of male ears to female WBFs that underlies male phonotaxis (Fig 1A) ^5–8^. Sexually dimorphic tuning frequencies are shaped by highly dimorphic auditory processes within the JO.

Interestingly, and uniquely among insect species, the tuning frequencies of mosquito ears are also in part modulated by an auditory efferent system which descends from the Central Nervous System (CNS) to innervate distinct regions within the JO, with these innervation patterns being sexually dimorphic ^9^. Beyond its’ role in modulating ear tuning frequencies, the auditory efferent system of mosquitoes also controls the onset of self-sustained oscillations (SOs) in male ears, an auditory phenotype characterized by large amplitude, monofrequent oscillations of the flagellum that reflects enhanced auditory amplification. SOs are not found in female ears ^10^.

Although sexual dimorphisms in *Ae. aegypti* acoustic communication systems have been characterized before, the molecular mechanisms shaping these various dimorphisms remain poorly understood, especially regarding the formation of female-specific features ^11–13^.

Sex-specific alternative splicing of the sex determination gene *doublesex* (*dsx*) produces sex-specific Dsx protein isoforms that differentially regulate somatic cell differentiation between sexes. The actions of different Dsx protein isoforms in turn determine the development of sexually dimorphic morphological structures that support sex-specific physiology and behaviors ^14^. Prior work in *Anopheles gambiae* (*An. gambiae*) mosquitoes has found that loss of the female isoform of *dsx* (*dsxF*) alters female anatomy and behaviors ^15,16^. However, how DsxF shapes female anatomy and behaviors remains unclear.

Here, we found that the loss of DsxF in female *Ae. aegypti* mosquitoes resulted in a flightless phenotype. By combining analyses of newly collected bulk RNA-seq data from mosquito thoraces with a previously published single-nucleus RNA-sequencing (snRNA-seq) dataset ^17^, we identified the complete loss of expression of a female-specific myosin gene, *myo-fem*, in *dsxF* female thoraces, which explains their flightless phenotype ^18^. We provide molecular evidence confirming that DsxF promotes *myo-fem*’s expression through binding to multiple *dsx* motifs within the *myo-fem*’s enhancer region.

Next, we profiled the hearing system of *dsxF* mutant females. We consistently observed intermediate phenotypes that are in between those of males and females throughout our anatomical and functional investigations. Anatomically, we observed that *dsxF* mutant female ears contained more plumose flagella and larger JOs than that of the females, as well as a partially masculinised auditory efferent system. Functionally, *dsxF* mutant females exhibited increased ear tuning frequencies compared to that of the control females, though not to the extent of the males. *dsxF* mutant females also did not gain male-specific SOs. Finally, analysis of newly collected bulk RNA-seq data from *dsxF* mutant female ears found significant upregulation of essential hearing genes compared to that of the control females. The enhancer regions of these genes bear *dsx* motifs, which are bound by DsxF, supporting DsxF’s role in directly inhibiting the expression of these genes.

We propose that DsxF plays distinct transcriptional roles in female *Ae. aegypti* thoraces and ears. In the thorax, it promotes the expression of *myo-fem* to enable female flight; on the other hand, in the ear, it inhibits the expression of key hearing genes to specifically tune female ears to lower frequencies compared to male ears. Lifting the inhibition of DsxF in female ears is not sufficient for complete masculinization however, indicating the presence of key male-specific transcription factors that drive greater extent of masculinisation in males.

## Results

### DsxF activates myo-fem’s expression via direct binding to support female flight

To investigate if the female-specific isoform of *dsx*, DsxF is involved in establishing female WBFs, we generated a *dsxF* mutant allele in *Ae. aegypti*, which contains an insertion of *3xP3-tdTomato* at the female-specific exon 5b (Figs 1B and 1C). We found that by disrupting DsxF, homozygous negative *dsxF* females (hereafter denoted *dsxF* females) develop intersex morphological structures (Figs S1A and 1B), including a male-like plumose flagellum and claspers (Fig S1A).

We further found that *dsxF* females in *Ae. aegypti* are flightless (Fig 1D, Supp Video 1). We hypothesized that the flightless phenotype of *dsxF* females could be due to disruption of flight-muscle gene expression. To test this, we dissected and submitted thoraces for bulk RNA-sequencing (Fig 1E). Our analysis of this bulk RNA-seq data confirmed sex-specific alternative splicing of *dsx* in male and female thoraces, suggesting differential roles of sex-specific Dsx isoforms in regulating gene expression in the thorax (Fig 1F).

To identify female-specific flight muscle genes under the transcriptional activation of DsxF, we looked for genes significantly upregulated (log2FoldChange>0, padj<0.1) in control females compared to both *dsxF* females and control males, which lack DsxF (Figs S2A and S2B). We found that AAEL005656 (*myo-fem*), a myosin gene known to be essential for female flight ^18^, is exclusively expressed in female, but not in control male and *dsxF* female, thoraces, suggesting that DsxF is required to activate *myo-fem* expression (Fig 1G). To support this, we utilized a published *Ae. aegypti* snRNA-seq atlas to look for potential overlap in *dsx* and *myo-fem* expression ^17^. We observed exclusive *myo-fem* expression in the female-specific muscle cell cluster, with *myo-fem* exhibiting strong overlap in expression with *dsx* (Figs 1H, 1I and S2C).

To investigate if DsxF could directly promote *myo-fem* expression, we searched for potential Dsx binding sites in the 1000bp promoter region upstream of *myo-fem*’s translational start site (TSS). Dsx proteins are zinc-finger transcription factors belonging to the Doublesex and Mab-3 Related Transcription factor (DMRT) family which share a highly conserved DM (*Doublesex and Mab-3*) DNA-binding domain that binds to DNA motifs with a consensus pseudopalindromic DNA element ^19^. We performed motif scanning using the well-characterized, highly conserved *Drosophila melanogaster* (*D. melanogaster*) Dsx*-*binding motif ^19^ and identified four potential Dsx binding-sites in the *myo-fem* enhancer region (Fig 1J). To examine if DsxF could directly bind to these sites, we performed Electrophoretic Mobility Shift Assay (EMSA) to assess interactions between the DsxF protein and four 100bp DNA sequences derived from the *myo-fem* enhancer region, each containing a centered Dsx-binding motif with native flanking sequences (Figs 1K and S2D). DsxF showed binding activity to all four *myo-fem* enhancer fragments, with these interactions lost when using truncated DsxF without a DM DNA-binding domain.

To summarize, we observed that DsxF binds to and promotes the expression of *myo-fem* in female thoraces to support female flight.

### DsxF shapes female-specific ear neuroanatomy

We found that disrupting DsxF partially masculinizes the external morphology of *dsxF* females, including the development of shorter flagellae (Tables S1 and S2; pairwise t-test with BH corrections; p=5.76 x 10^-4^) but enlarged pedicels (which house the JO) compared to control females (Figs S1A and S1B). These morphological abnormalities were not limited to external appearance. *dsxF* female JOs were significantly larger than control females, but remained significantly smaller than control males (Tables S3 and S4; pairwise t-test with BH corrections; p=2.98 x 10^-8^ and p=3.86 x 10^-6^ for comparisons with control females and males), likely reflecting an increase in the number of JO neurons with the loss of DsxF (Figs 2A and 2B).

**Figure 2:**
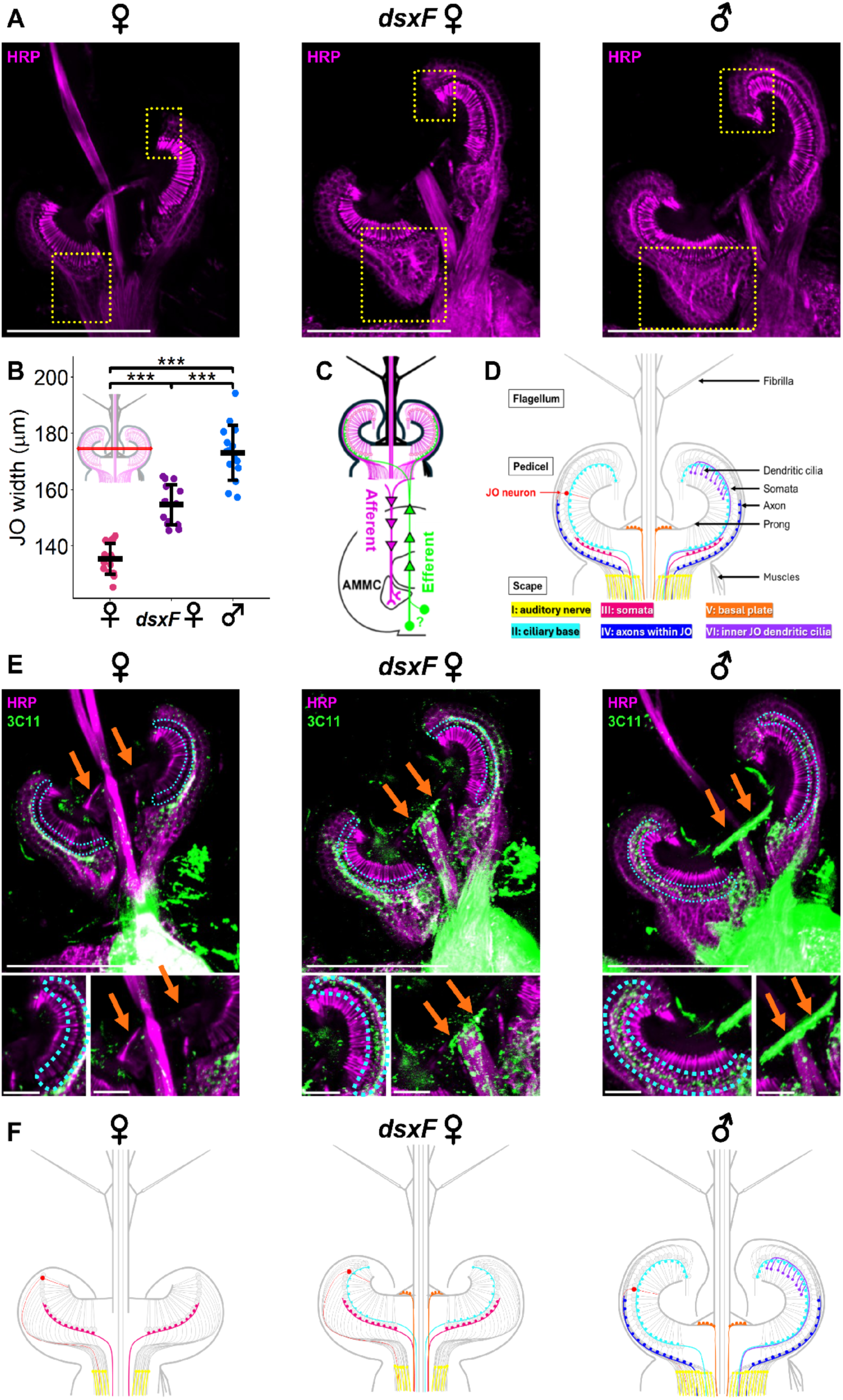
DsxF shapes female-specific ear neuroanatomy (A) JO sections of control females, *dsxF* females and control males stained with anti-HRP (magenta) to visualize JO neurons. JO regions with qualitative differences between genotypes boxed in yellow. Scale bar, 100μm; (B) Quantification of differences in JO width between genotypes, Individual points represent JO widths obtained from individual JOs. Solid lines represent mean and standard deviation. ***, p<0.001; pairwise t-tests with BH correction. See Tables S3 and S4 for measured values and statistical comparisons. Sample sizes: 15 control females; 14 *dsxF females*; 15 control males. (C) Schematic of mosquito auditory afferent and efferent systems. The auditory afferent system consists of JO neurons sending axonal projections to the Antennal Mechanosensory and Motor Center (AMMC) in the brain (arrowheads indicate direction of information flow). The auditory efferent system has unclear origin but is composed of efferent neurons with axonal projections that migrate and extend into the JO, which innervate and synapse onto different JO regions. (D) Schematic of classification of auditory efferent presynaptic terminal types based on innervation pattern in the mosquito JO. Adapted from ^11^. (E) (top) Auditory efferent presynaptic terminals within the JOs of control female, *dsxF* female and control males visualized using immunofluorescence. Maximum intensity projections of JO sections stained with anti-SYNORF1 (3C11, green) and anti-HRP (magenta). Dotted cyan outline = type II terminals (ciliary base), orange arrows = type V terminals (basal plate). Scale bar, 100μm. (bottom) Close-up images of type II (dotted cyan outline, ciliary base) and type V (orange arrows, basal plate) presynaptic terminals in the JO across genotypes. See Fig S3A for anti-SYNORF1 (3C11) channel images. (F) Schematic of differences in JO neuroanatomy between groups, focusing on differences for both JO size and auditory efferent innervation.

Mosquito ears adopt a bidirectional signaling system that modulates auditory signal transduction and transmission in a highly synchronized fashion. In addition to relaying auditory signal from the ears to the brain (“afferent signaling”), mosquito ears contain an auditory efferent system that transmits signals from the central nervous system (CNS) to the ears (“efferent signaling”) (Fig 2C). The mosquito auditory efferent system is a unique feature not identified in other insect species but found in mammalian cochlea, where it plays an important role in controlling hearing sensitivity ^9,20^.

The auditory efferent system is composed of neurons descending from the CNS that innervate distinct regions of the JO, which in turn modulate JO neuron properties, and thus hearing function, *via* neurotransmitter release. Previous reports found sexual dimorphisms in the localization of efferent presynaptic terminals within the JO, reflecting dimorphisms in neuromodulation of JO function ^8,9^. To determine if DsxF is involved in establishing sexually dimorphic efferent presynaptic localization, we stained JO sections with anti-SYNORF1 (3C11) antibody and categorized efferent terminals based on their localization following previous classification (Figs 2D and 2E). We observed that whilst still retaining female-specific type III terminals, *dsxF* females gained male-specific type II and type V terminals (Figs 2E, 2F and S3A).

Taken together, DsxF shapes female-specific JO neuroanatomy by defining both JO neuron number and the auditory efferent innervation within female JOs.

### DsxF defines female-specific peripheral hearing function

Mosquito flagellar ears are highly sensitive mechanosensors ^6,7^. Flagellar deflections stretch-activate mechanotransducers in JO neurons that convert mechanical forces into electrical signals relayed to the brain. Beyond transducing and transmitting auditory information to the brain, JO neurons also actively inject energy into setting the flagellar motion, which determines the frequency sensitivity and selectivity of mosquito ears ^10^. Male *Ae. aegypti* ears are tuned to higher frequencies of sound than female ears, reflecting enhanced amplification and auditory sensitivity across distinct frequency ranges tailored to sex-specific hearing needs ^12^. Male ears are tuned to female WBFs, facilitating male phonotaxis.

Given the important role of DsxF in defining female-specific JO neuroanatomy, we next asked if the altered JO neuroanatomy of *dsxF* females, including their potentially increased JO neuron counts and altered efferent presynaptic localization, could alter their hearing function. We used a laser Doppler vibrometer to measure mechanical vibrations of the mosquito flagellum whilst simultaneously measuring mechanically-evoked compound action potentials (CAPs) *via* a recording electrode inserted at the antennal nerve (Fig 3A). By stimulating mosquito ears with 1-1000Hz frequency sweeps *via* electrostatic actuation (Figs 3A and S4A), we found that *dsxF* females displayed intermediate mechanical and electrical tuning frequencies in between those of control females and males (Tables S5, S6 and S7; pairwise t-tests with BH correction; p=8.64×10^-11^ and p=3.16×10^-4^ for mechanical tuning; p=7.00×10^-5^ and p=2.96×10^-7^ for electrical tuning; Fig 3B).

**Figure 3:**
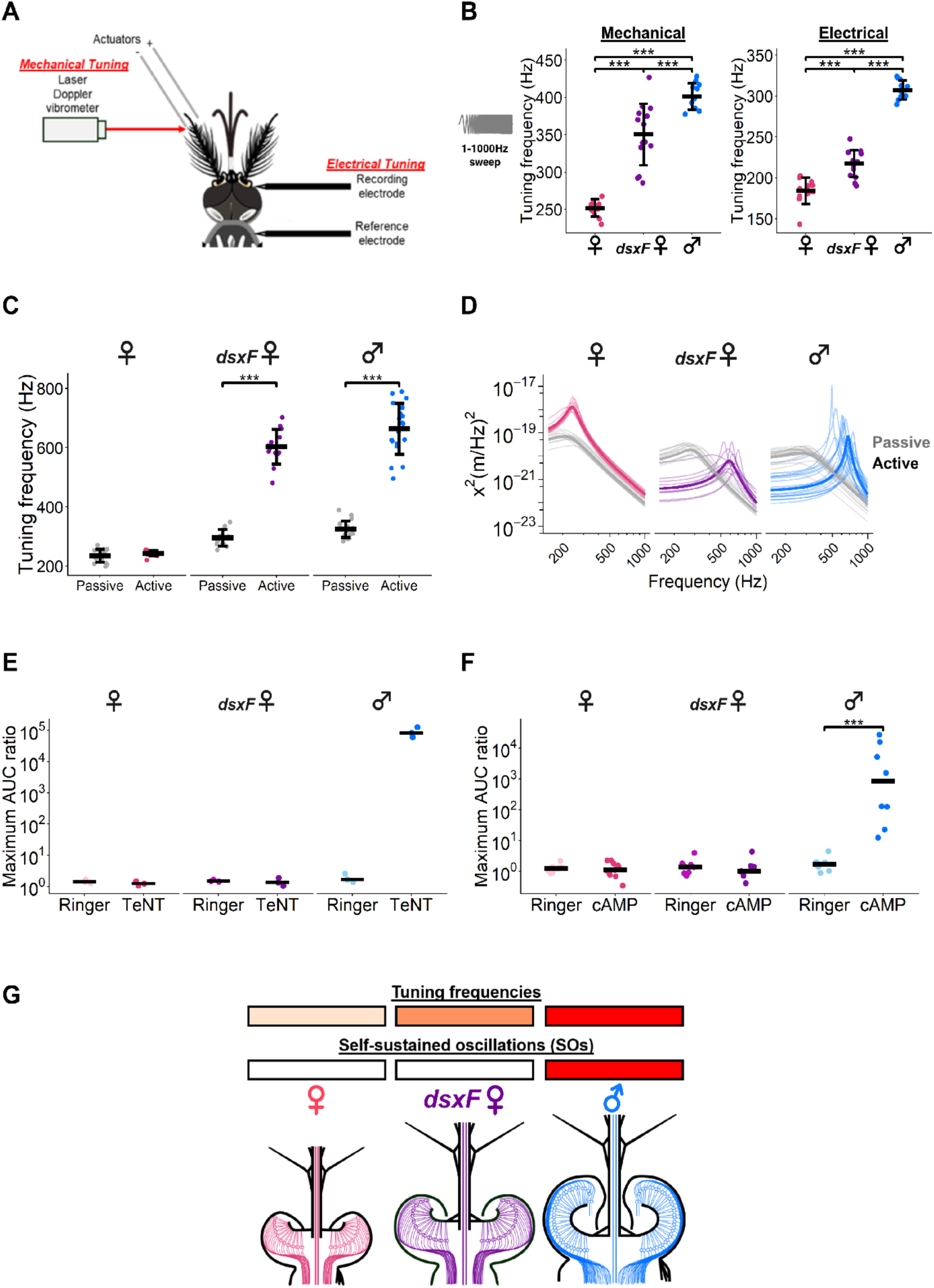
DsxF defines female-specific peripheral hearing function (A) Schematic of combined laser Doppler vibrometry and electrophysiology assay. Adapted from ^11^. (B) Active, sweep stimulated mechanical (left) and electrical (right) tuning frequencies of control females, *dsxF* females and control males. Individual points represent measurements from individual mosquitoes. Solid lines represent means and standard deviations. ***, p<0.001, pairwise t-tests with BH correction. See Tables S5, S6 and S7 for measured values and statistical comparisons. Sample sizes: 19 control females; 18 *dsxF* females; 17 control males. (C) Active and passive unstimulated mechanical tuning frequencies of control females, *dsxF* females and control males. Individual points represent measurements from individual mosquitoes. Solid lines represent means and standard deviations. ***, p<0.001, pairwise t-tests with BH correction. See Tables S8 and S9 for measured values and statistical comparisons. Sample sizes: 15 control females; 13 *dsxF* females; 19 control males. (D) Active and passive state unstimulated mechanical fluctuations of control female, *dsxF* female and control male flagellar ears. Thin lines represent fits for individual mosquitoes, thick lines represent fits to median parameter values. Grey lines represent passive state mechanical fluctuations and colored lines represent active state mechanical fluctuations. (E) Maximum ratio of AUC for vibrometry recordings taken before and after compound injection (Ringer or TeNT) for control females, *dsxF* females and control males. Individual points represent data from individual mosquitoes. Solid lines show medians. See Table S10 for maximum ratio of AUC following Ringer or TeNT injections. Sample sizes for Ringer and TeNT injections: 3/3 control females; 3/3 *dsxF* females; 3/3 control males. (F) Maximum ratio of AUC for vibrometry recordings taken before and after compound injection (Ringer or cAMP) for control females, *dsxF* females and control males. Individual points represent data from individual mosquitoes. Solid lines show medians. ***, p<0.001, Wilcoxon tests. See Tables S11 and S12 for maximum ratio of AUC following Ringer or cAMP injections and p values. Sample sizes for Ringer and cAMP injections: 7/10 control females; 8/7 *dsxF* females; 8/8 control males. (G) Schematic of differences in tuning frequencies between groups and absence/presence of self-sustained oscillations (SOs) in each genotype.

To examine if the increase in tuning frequencies of *dsxF* females compared to control females was due to active contributions from JO neurons, or a by-product of changes in ear structural and cuticular properties, we measured unstimulated flagellar motions in active and passive states. To reveal passive flagellar motions, we reversibly sedated mosquitoes using CO2. We observed higher passive (sedated) state mechanical tuning frequency of *dsxF* female ears compared to control females, indicating fundamental structural changes of female ears upon losing DsxF, matching the above anatomical and morphological observations. However, we still found a significant difference in the mechanical tuning frequencies of *dsxF* females between active and passive states, indicating that changes in tuning frequencies were still in part of active origin (Tables S8 and S9; t-test; p=4.16×10^-10^; Figs 3C and 3D).

Another signature of sexually dimorphic hearing function is the unique ability of male ears to show self-sustained oscillations (SOs), which are large, monofrequent flagellar vibrations with amplification gain hundreds to thousands of times above baseline, powered by motile properties of JO neurons ^10^. The onset and offset of SOs are modulated by the auditory efferent network *via* neurotransmitter release ^9,10^. Previous reports demonstrated that injection of tetanus toxin (TeNT), which blocks presynaptic release of neurotransmitters, could trigger the onset of SOs only in males ^10,11^].

Given that we observed partially masculinized JO neuroanatomy and hearing function in *dsxF* females compared to control females, we wondered if *dsxF* female ears also gained the ability to show male-unique SOs. By injecting TeNT into all three genotypes, we observed that only control males showed SOs (Table S10; Fig 3E).

Another method to induce SOs in males is by injecting 8-Br-cAMP, which triggers SOs by directly activating JO neurons ^11,21^. Consistent with TeNT injections, we observed SOs only in control males (Tables S11 and S12; Fig 3F). Therefore, we conclude that despite their partially masculinized ears, *dsxF* females did not gain the ability to show SOs, suggesting that other male-unique factors could be directing male SOs.

Taken together, DsxF is essential in determining female-specific tuning properties of the flagellar ears (Fig 3G).

### DsxF inhibits male-biased hearing gene expression in female ears

The intrinsic molecular machineries of JO neurons dictate JO neuron properties that collectively shape the functional properties of flagellar ears. We hypothesized that the altered hearing function phenotype of *dsxF* females could be due to altered expression of hearing genes. We first confirmed the presence of sex-specific alternative splicing of *dsx* in male and female *Ae. aegypti* pedicels using published bulk RNA-seq data ^12^, supporting our hypothesis that sex-specific transcriptional actions of Dsx proteins could differentially regulate hearing gene expression in *Ae. aegypti* JOs (Fig S5A).

We next dissected and submitted *dsxF* female pedicels for bulk RNA-sequencing (Fig 4A). We respectively identified 2591 and 2469 genes significantly up- and downregulated (log2FoldChange>0 or <0, padj<0.1) in *dsxF* female pedicels compared to control females (Figs 4B and S5B). GO enrichment analyses suggest that upregulated genes are related to microtubule-based process and cell motility, while downregulated genes are related to metabolism and cellular respiration (Fig S5C). The former is highly relevant in the context of mosquito hearing, as JO neurons themselves are composed of 9+0 axonemal microtubules on which cilium-specific motor complexes act to confer the ciliary motility necessary to power active hearing ^4,12^.

**Figure 4:**
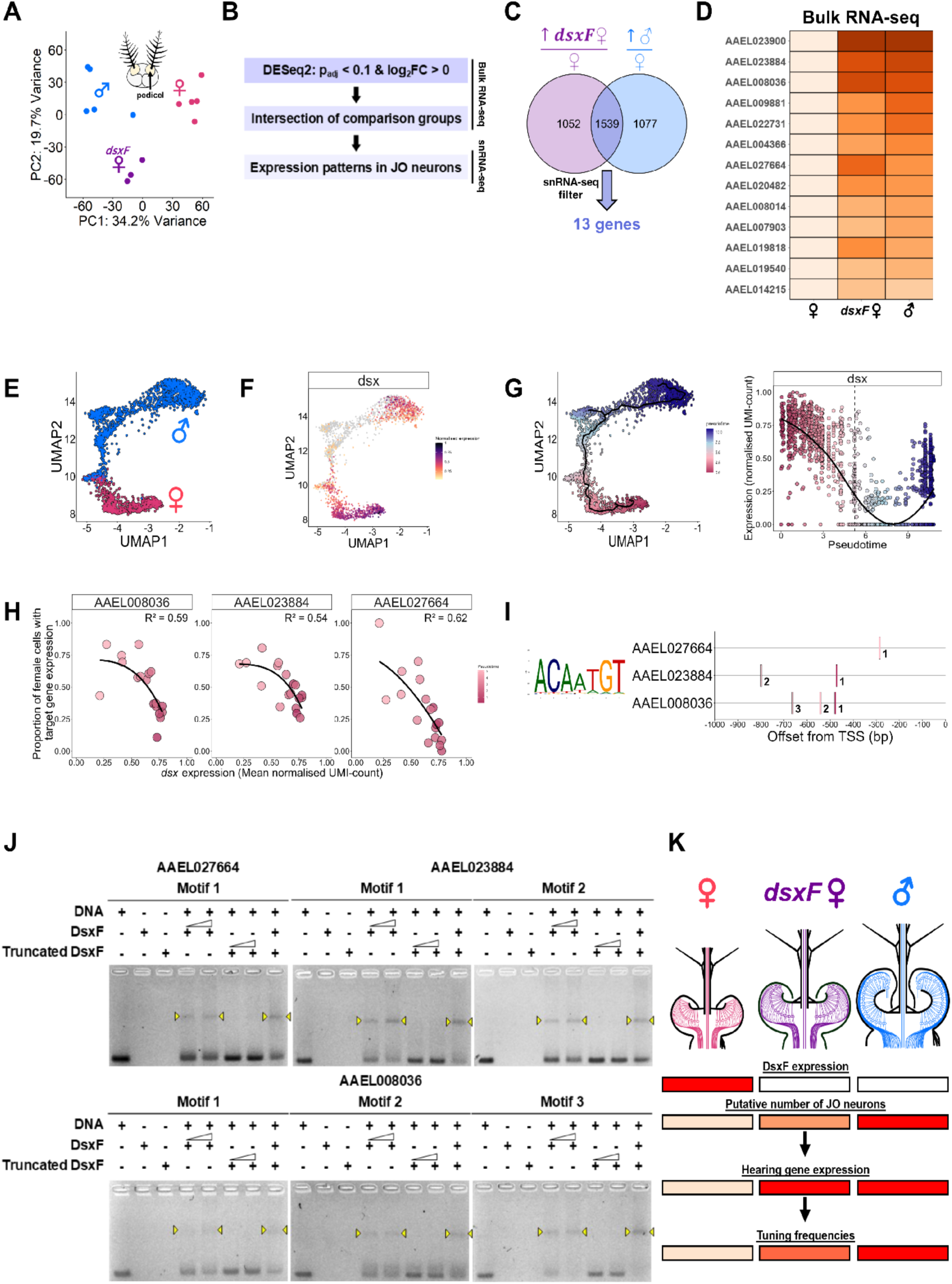
DsxF inhibits male-biased hearing gene expression in female ears (B) PCA of pedicel RNA-sequencing data for control females, *dsxF* females and control males. (C) RNA-seq data analysis process: first, differential gene expression analysis was conducted using DESeq2, followed by filtering for gene of interest by setting padj< 0.1 and log2FoldChange> 0; next, genes were selected based on intersections of comparisons group; finally, expression patterns in JO neurons were examined using published head snRNA-seq datasets where JO neurons were identified as *iav*-positive cells ^17^. (D) Venn diagram of pedicel genes upregulated in *dsxF* females and control males compared to control females. Of 1539 genes in the intersection, 13 were selected for further investigation based on expression patterns in JO neurons in published head snRNA-seq data. See also Fig S5D. (E) Heatmap of DESeq2 normalised counts of 13 genes identified from (**C**) in the pedicels of control females, *dsxF* females and control males from our pedicel bulk RNA-seq data. Color intensity reflects high/low expression level. See also Fig S5D. (F) JO neurons (identified as *iav*-positive cells) in published head snRNA-seq data ^17^. Male and female JO neurons identified based on sex of head samples. (G) Expression of *dsx* in JO neurons in published head snRNA-seq data ^17^. Color intensity represents normalized UMI-count. (H) (left) *pseudotime* plot indicating trajectory (path taken) when traversing from female to male JO neurons. Trajectory represented as *pseudotime* where time equals zero represents the start of the trajectory (distal-most female JO neurons) and maximum time represents end of trajectory (distal-most male JO neurons). Each nucleus colored based on *pseduotime* location on defined trajectory. (right) Changes in *dsx* expression level (in normalized UMI-count) along the defined trajectory. Each point represents normalised UMI-count in a single nucleus. (I) Proportion of female JO neurons expressing three different genes (AAEL008036, AAEL023884, AAEL027664) against mean normalised *dsx* UMI-count within each *pseudotime* bin. Individual points represent data grouped by *pseudotime* bins. Solid line represents repressive Hill function fitted to data points ^40^. R^2^ value calculated using the goodness-of-fit measure in the nls package. (J) Occurrence of Dsx-binding motif in the promoter region (1kbp upstream of translational start site) of AAEL008036, AAEL023884 and AAEL027664. (K) EMSA data showing protein-DNA interactions between full-length DsxF protein or truncated DsxF protein without DM domain and the 100 bp promoter fragments from genes that each bears a centered Dsx-binding motif. Interactions represented as band shifts were observed only between full-length DsxF protein and target DNA sequences. See Fig S6B for whole gel images. (L) Schematic linking changes in DsxF expression, JO neuron counts, and hearing gene expression to changes in tuning frequencies across genotypes.

We hypothesized that loss of DsxF could result in increased expression of hearing genes that in turn promote higher tuning frequencies. We therefore focused on genes significantly upregulated in both *dsxF* female and control male pedicels compared to control females (Fig 4C). By examining expression patterns of intersectional genes using published *Ae. aegypti* snRNA-seq data ^17^, we identified 13 genes with JO-biased expression (JO neurons defined as *inactive* expression > 0) (Figs 4C, 4D, S5D and S5E), with some known to be essential for hearing in *D. melanogaster* and mosquitoes, such as AAEL019818 (*nompC*), AAEL020482 (*inactive*), AAEL023900 (*Dhc36C*), AAEL022731 (*Dhc93AB*), AAEL009881 (*Dhc1*) and AAEL008036 (*Dnah3*) ^22–24^. Lack of DsxF thus resulted in a step-like increase in expression of essential hearing genes in *dsxF* female and control male pedicels compared to control females. The expression levels of these hearing genes were also not significantly upregulated in control male pedicels compared to *dsxF* females (Fig 4D).

Collectively, DsxF inhibits the expression of male-biased hearing genes in control female pedicels.

### DsxF transcriptionally inhibits expression of male-biased hearing genes via direct binding

To identify hearing genes under direct transcriptional inhibition of DsxF, we utilized published snRNA-seq data to examine expression levels of these hearing genes in male and female JO neurons at the single-cell level ^17^. We observed that JO neurons form a continuous, elongated cluster in which male and female cells are largely delineated from each other, reflecting sex-specific gene expression in JO neurons with transcriptional continuum when traversing between the sexes (Fig 4E).

We next examined *dsx* expression in JO neurons and observed that almost all female JO neurons are *dsx* positive, while *dsx* expression in male JO neurons is restricted to a fraction of distal-most JO neurons, indicating that DsxF has an important and widespread role in regulating gene expression in female JO neurons (Fig 4F). We next adopted *pseudotime* analysis to quantify changes in *dsx* expression along a continuous trajectory (Fig 4G). By anchoring on the distal-most population of female JO neurons and traversing along the trajectory, we observed a reduction in *dsx* expression during the transition from female to male JO neurons that gradually recovers when approaching the distal-most population of male JO neurons (Fig 4G).

We next adopted *pseudotime* analysis to directly correlate changes in *dsx* expression level with the proportion of JO neurons expressing candidate hearing genes, along the continuum of female and male JO neurons (Fig S6A). To identify hearing genes that are transcriptionally inhibited by DsxF, we applied a repressive Hill function to describe the relationship between *dsx* expression level and the proportion of female JO neurons expressing a target gene within *pseudotime* bins. We found that as the mean *dsx* expression level in female JO neurons within each *pseudotime* bin increases, the proportion of female JO neurons within these bins expressing three specific dynein genes, AAEL008036 (*Dnah3*), AAEL023884 (*Dhc98D*) and AAEL027664 (*Dnah9*), decreases (Figs 4H and S6A), supporting that DsxF inhibits the expression of these male-biased hearing genes in female JO neurons.

To examine if DsxF could directly bind to the enhancer region of these genes to inhibit their expression, we conducted motif scanning and identified Dsx-binding sites in the enhancer region of all three genes (Fig 4I). Our EMSA results indicate binding of DsxF to all predicted binding sites, with these protein-DNA interactions lost when using truncated DsxF lacking the zinc-finger DM DNA-binding domain (Figs 4J and S6B).

By acting as a transcriptional inhibitor, DsxF thus binds to and inhibits the expression of male-biased dynein genes that could power the JO neuron motility necessary for achieving the higher ear tuning frequencies of males (Fig 4K).

## Discussion

Here, we identified important roles of DsxF in modulating both flight and hearing systems of female *Ae. aegypti* mosquitoes. The loss of DsxF renders *dsxF* females flightless due to the lack of expression of a female-specific myosin gene (*myo-fem*) in the thorax, consistent with previous reports ^18^. Our work expands on prior findings by highlighting DsxF as an upstream transcriptional activator of *myo-fem* (Fig 1K).

Our *Ae. aegypti* findings are distinct from work in *An. gambiae*, where loss of DsxF instead increases *dsxF* mutant female WBFs compared to control females, though the effector genes responsible remain unclear ^15^. These interspecific differences could stem from differences in DsxF-mediated transcriptional programs regulating expression of flight muscle genes, different genes involved in flight or the interplay between flight muscle genes ^15^.

We observed masculinization of morphological characteristics in *dsxF* females compared to control females. The flagellar ears of *dsxF* females contain more plumose flagella and larger pedicels than control females. The pedicels of *dsxF* females also potentially house more JO neurons than control females (Fig 2). However, the degree of masculinization in *dsxF* females is incomplete ^25^. The formation of sexually dimorphic flagellar ears therefore relies on DsxF to inhibit extensive morphogenesis and neurogenesis in females during development and potentially other male-specific factors, such as the transcriptionally opposing actions of DsxM to drive greater extents of cell proliferation and differentiation in males. *doublesex* is the second sex determination transcription factor found to be involved in shaping sexually dimorphic auditory efferent networks in *Ae. aegypti* mosquitoes, in addition to *fruitless* ^11^. However, how DsxF inhibits the formation of type II and type V presynaptic terminals within female JO remains unknown. Future developmental studies could focus on elucidating the DsxF-mediated developmental programs that shape dimorphic flagellar ears and auditory efferent networks.

Functionally, we found that *dsxF* females gained higher active state tuning frequencies compared to control females (Fig 3G). We associated this increase with: 1) a likely increased number of JO neurons and 2) the lack of inhibition of male-biased hearing gene expression (Fig 4K). However, we still observed substantially higher tuning frequencies for male ears compared to *dsxF* females, as well as male-specific SOs (Fig 3G). These differences could be attributed to: 1) retention of non DsxF-dependent female-unique hearing gene expression, 2) lack of/insufficient expression of male-biased hearing genes, or 3) lack of male-specific neuronal populations, in *dsxF* female ears. Future work combining mutagenesis (such as generating DsxM mutants) with high-resolution molecular, functional and anatomical investigations could help shed light on these issues.

Our findings in *Ae. aegypti* expands on prior knowledge learned from *An. gambiae*; by integrating our bulk transcriptomic datasets with published *Ae. aegypti* snRNA-seq data and complementing our computational analyses with biochemical evidence, we present unprecedented insights into how DsxF transcriptionally inhibits male-biased gene expression in female ears.

In summary, we identified transcriptionally opposing roles of DsxF in two distinct female tissues: as an activator in the thorax but an inhibitor in the ear. Our work highlights the important roles of DsxF in establishing female-unique properties across sexually dimorphic sensory (hearing) and motor (flight) systems to coordinate the acoustic communication systems of *Ae. aegypti* mosquitoes.

## Resource availability Lead contact

Requests for further information and resources should be directed to, and will be fulfilled by, the lead contact, Matthew P Su.

## Materials availability

Requests for the *dsxF* mutant will be fulfilled pending scientific review and a completed material transfer agreement.

## Data and code availability

- Data are publicly available at the paper’s Mendeley data repository (available upon publication).
- All original code is available *via* Mendeley data (available upon publication).
- Any additional information required to reanalyze the reported data is available from the lead contact upon request.

## Acknowledgments

We would like to thank Mika Nomoto, Yasuomi Tada, Akiko Akama, and Mikako Yamaguchi (Center for Gene Research, Nagoya University) for assistance with RNA-seq experiments at the Division for Medical Research Engineering. We thank Yixiao Zhang for technical support with mosquito rearing.

This study was funded by the following grants:

JST FOREST JPMJFR2147 (AK)

MEXT KAKENHI Grant-in-Aid for Scientific Research (B) JP26K01748 (AK)

MEXT KAKENHI Grant-in-Aid for Transformative Research Areas (A) “Materia-Mind” JP24H02200 (AK)

Human Frontier Science Program Organization RGP0033/2021 (AK)

MEXT KAKENHI Grant-in-Aid for Research Activity Start-up JP22K15159 (MPS) Nagoya University Tokai Pathways to Global Excellence 0121an0002 (MPS) JSPS Invitational Fellowships for Research in Japan (Short-term) S22091 (DFE)

International Principal Investigator (PI) Invitation Program, Nagoya University, Japan (DFE)

JSPS PhD 2524KJ1285 (YML)

UCSD Discretionary funds (OSA)

## Author contributions

Conceptualization YML, DFE, MPS, AK

Data Curation YML, MPS, AK

Formal Analysis YML, MPS

Funding Acquisition YML, DFE, OSA, MPS, AK

Investigation YML, WWL, JXY, TTL, YYJX, ML, MPS

Methodology YML, WWL, JXY, YT, DFE, ML, OSA, MPS

Project Administration YML, MPS, AK

Resources YML, DFE, OSA, MPS, AK

Software YML, MPS

Supervision OSA, MPS, AK

Validation YML, MPS

Visualization YML, MPS

Writing – Original Draft YML

Writing – Review and Editing YML, DFE, ML, OSA, MPS, AK

## Methods

### Animals used in this study

All mosquitoes tested were 5-7 day old adults. Pupae were sex-separated to ensure adults were virgin, with glucose water provided throughout the adult stage. Tested females were not blood fed. Mosquito lines used in this study were wild-type *Ae. aegypti* (Liverpool strain) and *Ae. aegypti dsxF* mutants (Liverpool background).

### Mosquito rearing

All mosquitoes were reared at 28°C and 60-70% relative humidity in 12 h:12 h Light:Dark (LD) conditions. Larvae were provided with fish food and adults were provided with 10% glucose water. Blood feeding utilised a membrane feeding system (Orinno Technology Pte Ltd., Singapore).

### Experiment entrainment paradigm

Adult mosquitoes were entrained for two days under 12:12 LD conditions with constant access to 10% glucose water. Entrained mosquitoes were tested starting on the third entrainment day between Zeitgeber Time (ZT) 11-ZT13, corresponding to the “dusk” phase. All mosquitoes tested were 5-7 days old. All experiments were conducted in a temperature-controlled room or incubator set at 22°C (±1.5°C).

### Mutagenesis

The knock in plasmid was cloned using Gibson enzymatic assembly ^26^. The plasmid (Addgene ID: 117221) was used as a template to insert the following fragments: (1) left and right homology arms of ∼1 kb in length, which are complementary to the *Ae. aegypti dsx* locus immediately adjacent to the 5’ and 3’ ends of the *dsx* gRNA cut site, respectively; (2) a U6 promoter with a 20-base *dsx* gRNA sequence, a 76-base gRNA scaffold and the 3’UTR region of the U6 snRNA; (3) a 3xP3-tdTomato-SV40 fragment. All DNA fragments were synthesized by GenScript.

The generated plasmid was transformed into Zymo JM109 chemically competent *E. coli* (Zymo Research, Cat. #T3005), amplified, isolated (Zymo Research, Zyppy plasmid miniprep kit, Cat. #D4036), and Sanger sequenced. The final plasmid was maxi-Xprepped (Zymo Research, ZymoPURE II Plasmid Maxiprep kit, Cat. #D4202) and sequenced using Oxford Nanopore Sequencing at Primordium Labs (https://www.primordiumlabs.com).

The *dsxF* transgenic strain was generated by microinjecting preblastoderm stage embryos (0.5–1 hr old) with a mixture of the donor plasmid (100 ng/µl), synthetic gRNA (40 ng/µl) and Cas9 protein (100 ng/µl). Embryonic collection, microinjection, transgenic line generation, and rearing were performed following previously established procedures^27^.

### Video recording of flight behavior

Cages of five control females and five *dsxF* mutant females were entrained as described above. Flight activity was recorded using a video camera (GoPro) on the third day between ZT11.5-ZT12 at 22°C (±1.5°C).

### Images of mosquito anatomy

Mosquitoes were sedated on ice and positioned for imaging using forceps. A camera (SONY α7S) was used to image mosquito heads and genitalia.

### Quantification of flagellar length

Mosquitoes were sedated on ice and their right flagellae removed. Samples were mounted on slide glasses and imaged using a confocal microscope (FV3000) with a 2x lens (UPLSAPO2x) at 2-fold magnification. Sample lengths were measured using ImageJ (version 1.53q, National Institutes of Health, RRID: SCR_003070).

Flagellar length was measured as the length of the flagellum from the tip of the first flagellomere to the end of the 13th flagellomere.

Sample sizes: 45 control females; 24 *dsxF* females; 20 control males.

### Sample preparation for RNA-sequencing

5-6 days old mosquitoes were collected at ZT12 and flash frozen in liquid nitrogen. For thorax samples, control females, *dsxF* females and control males were dissected. For head and pedicel samples, only *dsxF* females were dissected.

Mosquito thoraces, heads and pedicels were dissected in RNAiso Plus (9109, Takara Bio Inc) on ice, and the tissues were homogenized. Homogenized samples were added with RNAiso Plus until the final volume reached 1 mL. Samples were incubated at room temperature for 15 minutes, followed by the addition of 200 µL chloroform (Kanto Chemical Co., Inc.). The samples were mixed well by inverting the tubes several times followed by incubation at room temperature for 15 minutes.

Samples were centrifuged at 12,000g at 4°C for 15 minutes and the supernatant was transferred carefully to fresh tubes. 0.5 mL ice-cold 2-propanol (Sigma) was added into each tube followed by inverting the tubes several times to mix the samples well. The tubes were kept in −20°C for 30 minutes, followed by centrifugation at 12,000g at 4°C for 15 minutes. Supernatant was discarded and RNA pellets were washed twice with 1 mL 75% ethanol (Sigma). RNA pellets were air-dried followed by dissolving the pellets using DEPC water (Nacalai Tesque). A Nanodrop (ThermoFisher Scientific) was used to assess RNA quality for each sample.

For all thoraces and *dsxF* female heads and pedicels, three biological repeats for each group were submitted to the Center for Gene Research at Nagoya University for library preparation and sequencing.

### Transcriptomics: Library preparation, read alignment and differential expression analysis

The *Ae. aegypti* L5 genome fasta file, available *via* VectorBase (68^th^ release), was used for read-mapping ^28^. Single-end reads were mapped to this genome using the Rsubread package. Using the *Ae. aegypti* GFF file, available *via* VectorBase (68^th^ release), gene counts of aligned reads were obtained using Rsubread’s featureCounts function.

Differential expression analysis was conducted using the DESeq2 package ^29^ in R with a fold-change threshold of 1 and FDR cutoff <0.1. Pedicel analyses were conducted by including previously published pedicel datasets of wild-type male and female *Ae. aegypti* mosquitoes ^12^.

### *dsx* splicing analysis

DEXSeq was used for differential splicing analysis ^30^. A flattened GFF file was created using the aforementioned *Ae. aegypti* GFF file (68^th^ release, VectorBase) in DEXSeq. BAM files of aligned reads of *Ae. aegypti* male and female thoraces and pedicels were mapped to this annotation. Exon usage counts for *dsx* (AAEL009114) were calculated using RSubread’s featureCounts function ^31^.

### Examination of gene expression patterns in thorax using published snRNA-seq data

Expression patterns of target genes were plotted using the Monocle3 ^32^ package in R, based on published adult *Ae. aegypti* thorax snRNA-seq data ^17^. Male and female cells were identified based on sample sex.

### Motif identification

Promoter sequences (1kbp upstream of translational start site) were retrieved using the biomaRt::getSequence function in R ^33^. The *dsx* motif (of *D. melanogaster*) was downloaded from JASPAR ^34^. Occurrences and positions of the *dsx* motif were determined using MEME Suite’s FIMO algorithm with default parameters ^35,36^.

### Protein purification

Protein-coding sequences for full-length DsxF protein and truncated DsxF protein lacking DM domain were fused with a 6xHis-MBP-tag at the N-terminus and cloned into a p15TV-L plasmid (Addgene Plasmid #26093). Plasmids were transformed into Rosetta-gami 2 (DE3) (Sigma-Aldrich) to induce protein expression. Bacterial cells were grown in LB medium-supplemented with 100 μM ZnSO4 at 37°C and 180rpm until the optical density OD600 reached 0.5-0.8. Protein expression was induced by adding 100 μM of IPTG (Sigma-Aldrich) and overnight shaking at 18 °C.

After induction, cells were harvested and resuspended in HEPES and glycerol (final concentration of 100mM HEPES (pH 7.5), 10% glycerol). Cells were then lysed by sonication, and the lysate was centrifuged at 10,000rpm for 30 min at 4 °C. Supernatant was transferred to a fresh tube and Ni-NTA resin (Profinity™ IMAC Resin, Ni-charged) pre-equilibrated with extraction buffer (100 mM HEPES (pH 7.5), 150 mM NaCl, 100 μM ZnSO4 and 10% glycerol) was added into the supernatant. After 30 minutes of pull-down using a rotor at 4 °C, the beads were collected by centrifugation at 4,000rpm for 1 min. Supernatant was discarded and beads transferred to an Eppendorf before washing three times with a wash buffer (extraction buffer + 30 mM imidazole).

The protein was eluted in elution buffer (extraction buffer + 150 mM imidazole). Desalting and buffer exchange (20 mM HEPES (pH 7.5), 30 mM NaCl, 100 μM ZnSO4) was performed using HiTrap™ Desalting columns with Sephadex G-25 resin. Finally, protein was aliquoted and stored at −80 °C.

### Electrophoretic Mobility Shift Assay

DNA probes for Electrophoretic Mobility Shift Assay (EMSA) were labelled with Fluorescein-5-maleimide (Tokyo Chemical Industry) using 5′ EndTag DNA/RNA Labeling Kit (Vector Labs). DNA binding reaction contained 350 ng of labelled DNA, 3.25 ug of protein, 10xbinding buffer (1x binding buffer: 10 mM (NH4)2SO4, 0.2% Tween20, 5% glycerol, 1 mM DTT and 1 ug BSA) and exchange buffer in final volume of 20uL.

The reactions were incubated in darkness at room temperature for 1 h, followed by gel electrophoresis using 1.5% agarose gel (in 0.5xTB). 3 uL of 30% glycerol was added right before gel-loading. Gel electrophoresis was performed at 150V for 20 min and the resulting gel was viewed using a gel-viewer (Atto).

### JO immunofluorescence

Mosquitoes were sedated on ice and their heads were removed. Samples were fixed for 1 h in 4% paraformaldehyde (PFA) in Phosphate-buffered saline (PBS) with 0.25% Triton X-100 (PBT) and embedded in 10% agarose solution (Agarose S, 318-01195). All samples were left at 4°C for 10 mins, then fixed overnight in 4% PFA at 4°C.

Samples were washed in 100% methanol for 10 min, then transferred to PBS for 30 min. Blocks were sectioned using a vibratome (Leica VT1200S, 40 μm thickness), with sections then washed in 0.5% PBT three times at room temperature. Samples were blocked in 10% normal goat serum (NGS) (Vector Laboratories, Inc.)/0.5% PBT for 1 h and incubated at 4°C with primary antibodies in 10% NGS/0.5% PBT for three days.

After this incubation, samples were washed three times with 0.5% PBT then incubated for two days at 4°C with secondary antibodies in 10% NGS/0.5% PBT.

After this, samples were washed three times with 0.5% PBT followed by one wash with PBS. Sections were mounted on slides and imaged using a laser-scanning confocal microscope (FV3000, Olympus) with a 20× air objective (UPlanSApo, NA = 0.75) lens.

Primary antibody: anti-SYNORF1 (3C11) monoclonal antibody (AB_528479, 1:30, Developmental Studies Hybridoma Bank (DSHB), University of Iowa).

Secondary antibodies: Alexa Fluor 488-conjugated anti-mouse IgG (A-11029, 1:300, ThermoFisher) and Cy3-conjugated goat anti-horseradish peroxidase (anti-HRP, RRID: AB_2338959, 1:150, Jackson Immuno Research).

### JO width measurements

Images of JO sections stained with HRP were used for measurements in ImageJ (version 1.53q)^37^. JO widths were only included in the analysis if the basal plate was clearly visible. The longest distance between the outer somata of both sides of the JO was measured as the JO width.

Sample sizes: 15 control females; 14 *dsxF* females; 15 control males.

### Preparation for laser Doppler vibrometry and electrophysiology

Mosquitoes were prepared following prior protocols ^12,13,21^. Sedated mosquitoes were glued to plastic rods with blue-light-curable glue (Norland Products Inc., 81). Following gluing, only the mosquito’s right flagellum was able to move. Prepared mosquitoes were tested in a temperature-controlled room (22 ± 1.5 °C), with the plastic rod inserted into a micromanipulator (MM-3, Narishige Instruments, Japan) on a vibration isolation table.

Vibrometry recordings used a laser Doppler vibrometer (Vibroflex, Polytec) in conjunction with the VibSoft software (Polytec). The laser Doppler vibrometer was focused on the first flagellomere from the tip of the flagellum.

For electrophysiology experiments, a tungsten electrode was inserted into the thorax to allow for charging to −30 V (relative to ground). Actuators were placed around the flagellum to allow for provision of electrostatic stimulation. A tungsten electrode inserted into the pedicel base enabled antennal nerve recordings. Electrophysiology and vibrometry recordings were collected simultaneously using the Spike2 software (ver 10.08, CED), with stimuli created in Spike2.

### Electrophysiology: Recordings

Recordings followed prior protocols ^12,13,21^. A calibrated force-step stimulus was first provided to calibrate flagellar displacements to approximately ± 2.5μm. Sets of sweep stimuli, including one-second-long chirps of linearly increasing (forward, 1 to 1000 Hz) or decreasing (backward, 1000 to 1 Hz) frequency, were used to stimulate flagellar ears. The flagellar displacement (100 kHz sampling rate) and antennal nerve responses (20 kHz sampling rate) were recorded in Spike2. Laser quality data (10 kHz sampling rate) was also recorded to facilitate downstream analyses.

### Electrophysiology: Sweep analysis

First, for flagellar mechanical tuning analyses, all data for the corresponding laser quality data was recorded as zero were excluded for further analyses. Channel processes were applied to the laser data as follows: a DC remove function (0.01s time constant); rectification; a smooth function (0.0005s time constant); a slope function (0.1s time constant). Processed data enabled identification of when the channel slope was equal to zero (time of maximum flagellar vibration). The stimulus frequency at this timepoint was used as the peak ear mechanical tuning frequency for that sweep. The average of consecutive phasic and anti-phasic sweeps of the same orientation was taken, with the median of all averages calculated as the peak mechanical tuning per mosquito.

For electrical tuning analyses, nerve responses to sequential phasic and anti-phasic stimuli were averaged to cancel out artefacts arising from electrostatic actuation. Nerve channel data then had the following processes applied: a DC remove function (0.01s time constant); rectification; a smooth function (0.0005s time constant); a slope function (0.1s time constant). Processed data enabled identification of when the channel slope was equal to zero (time of maximum antennal nerve response). The stimulus frequency at this timepoint was used as the peak ear mechanical tuning frequency. The average between sequential forward and backward sweeps was calculated, with the median of these averages calculated as the peak electrical tuning per mosquito.

Control male and female data was reanalysed from a previous publication, with recordings taken in identical conditions ^11^.

Sample sizes: 19 control females; 18 *dsxF* females; 17 control males.

### Laser Doppler vibrometry: Recordings

Unstimulated recordings of the free-fluctuating mosquito flagellum were recorded in both active and passive states. First, an active-state free-fluctuation recording was obtained, before mosquitoes were exposed to CO2 in a sealed chamber to induce sedation. An unstimulated recording of the flagellar ear was taken immediately after cessation of CO2 delivery.

### Laser Doppler vibrometry: Data analysis

Raw time-domain data between 1 and 1000Hz was Fast Fourier transformed using the VibSoft software (Polytec), with values below 125Hz excluded from further analyses due to noise.

A previously published forced damped oscillator function was fit to transformed data using the following formula in the lme4 package ^38^:

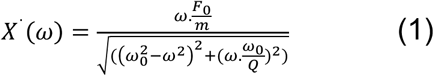

F0 = external force strength, m = flagellar apparent mass, ω = angular frequency, ω0 = natural angular frequency and Q = quality factor = mω0/γ (γ = damping constant).

Fit parameter estimation enabled calculation of the natural angular frequency (ω0) for each recording, and thus calculation of the flagellar ear mechanical tuning frequency, f0, based on f0= ω0/2π.

Control male and female data was reanalysed from a previous publication, with recordings taken in identical conditions ^11^.

Sample sizes: 15 control females; 13 d*sxF* females; 19 control males.

### TeNT and cyclic AMP (cAMP) thoracic injections for laser Doppler vibrometry

Mosquitoes were mounted as described above for vibrometry experiments. A free fluctuation recording was taken using the VibSoft software (Polytec) to act as a baseline prior to injection. Following this, compounds were injected into the mosquito and further recordings were made.

For TeNT injection, 2μM TeNT (T3194, Sigma) or Ringer’s solution was injected into the mosquito thorax. Recordings were made every 5 minutes for 60 minutes following injection.

For cAMP experiments, either 3mM 8-Bromoadenosine 3,5-cyclic monophosphate sodium salt (8-Br-cAMP; 76939-46-3, Sigma) or Ringer’s control ^8,21^ solution was injected into the mosquito thorax. Recordings were made 5, 10, 15 and 20 mins following injection. Recordings were analyzed as described above to extract the peak mechanical tuning frequency, enabling calculation of frequency change (ΔFrequency) at each recording timepoint compared to baseline. The largest ΔFrequency within an individual recording was taken as the maximum ΔFrequency.

Fluctuation power was calculated by first converting data to the displacement squared domain, then estimated using the AUC function in R. By dividing the AUC for each post-injection timepoint by the pre-injection baseline, the maximum AUC ratio following injection was calculated.

Control male and female data was reanalysed from a previous publication, with recordings taken in identical conditions ^11^.

Sample sizes for TeNT / Ringer injections: 3/3 control females; 3/3 dsxF females; 3/3 control males.

Sample sizes for 8-Br-cAMP / Ringer injections: 10/7 control females; 7/8 dsxF females; 8/8 control males.

### Gene Ontology enrichment analysis

A published modified GAF file ^11^ was used to run GO enrichment analysis using g:Profiler ^39^. This custom GMT file can be found using the following g:Profiler token: gp E7yL_EL9T_8t8. The SCS threshold algorithm was selected for significance testing with a significance threshold of 0.05 applied.

### Examination of gene expression patterns in JO neurons using published snRNA-seq data

Expression patterns of target genes were plotted using the Monocle3 package in R using published adult *Ae. aegypti* head snRNA-seq data ^17,32^. Male and female cells were identified based on sample sex. JO neurons were identified based on filtering for cells with *inactive* (AAEL020482) expression.

### Pseudotime analysis

*Pseudotime* analysis was performed using Monocle3 ^32^. Cells of interest (*inactive*-positive JO neurons) were first isolated from other head cells, and cells were ordered by setting distal-most female JO neurons as root cells (*pseudotime* equals zero) and the *pseudotime* spent traversing from distal-most female JO neurons to distal-most male JO neurons were computed. Changes in target gene expression levels along the computed *pseudotime* were then extracted for further analysis.

### Fitting of repressive Hill function

A repressive Hill function ^40^ was utilised to describe the relationship between *dsx* mean expression level in female JO neurons within each *pseudotime* bin and the proportion of female JO neurons expressing target gene within these bins, with the following formula:

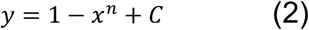

where x is the mean *dsx* expression level, y is the proportion of cells expressing each gene per *pseudotime* bin and n and C are constants.

Model fitting was performed using the nls function in R and fit quality was evaluated using the goodness-of-fit measure (R^2^). Fits with an R^2^ greater than 0.5 were selected for inclusion.

## Quantification and statistical analysis

P < 0.05 (prior to correction) was set as the significance level for all statistical testing. Two-sided statistical tests were used throughout.

Shapiro-Wilk tests of normality were first used to test for normality for all functional assay, flagellar length and JO width data. Flagellar length and JO width data were found to be normally distributed, so pairwise t-tests with BH corrections were used for statistical testing. Functional datasets, including sweep vibrometry/electrophysiology data, were all found to be normally distributed; as such, pairwise t-tests with BH corrections were used. Comparisons of unstimulated vibrometry recordings in active and passive states used t-tests, as unstimulated mechanical tuning frequencies were found to be normally distributed.

Vibrometry datasets for Ringer and cAMP injections were found to be non-normally distributed, and as such pairwise Wilcoxon tests with BH corrections were used.

