## Supplemental Information for "The sex determination gene *doublesex* is essential for female-specific flight and hearing properties in *Aedes aegypti* mosquitoes"

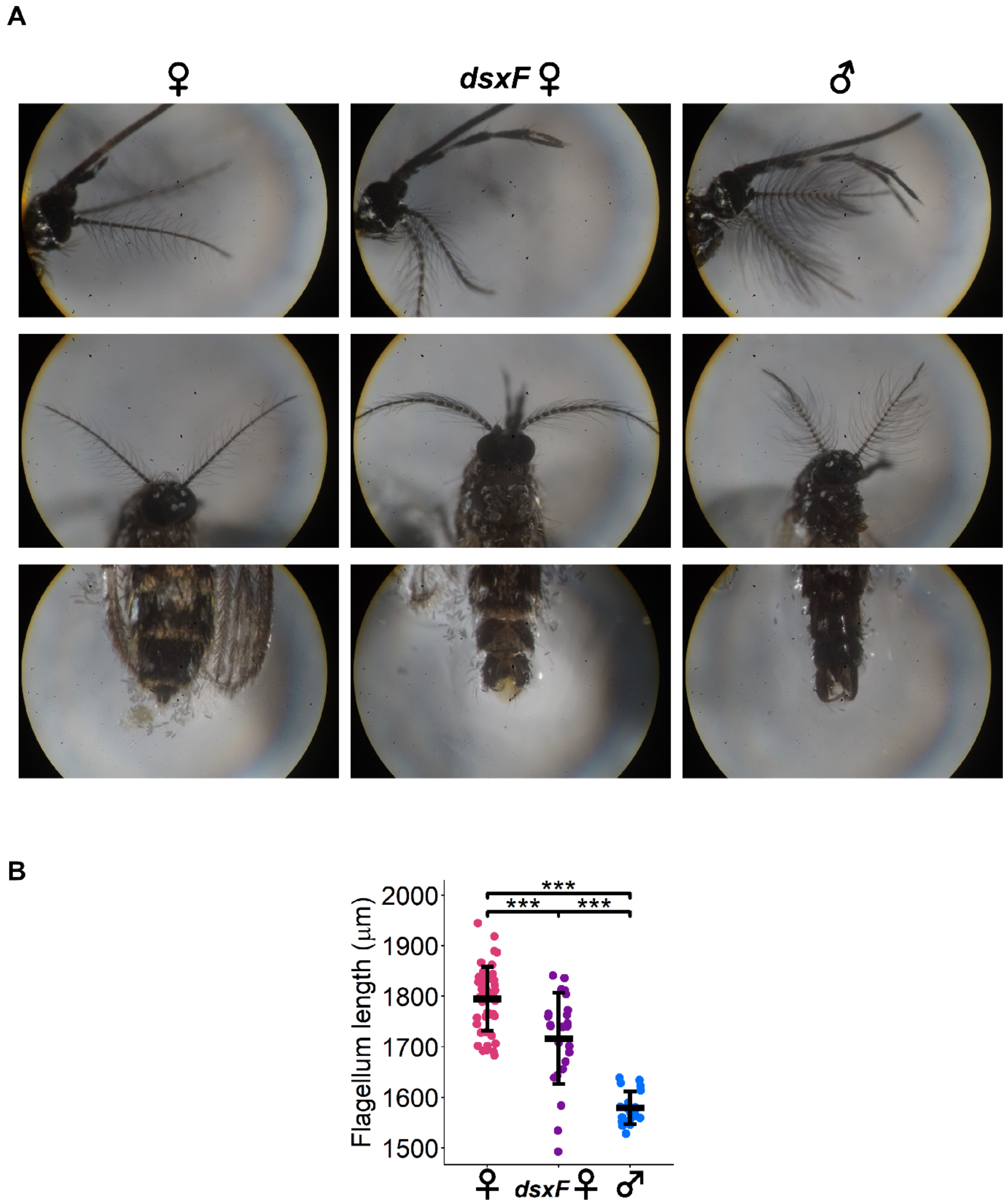

**Figure S1: *dsxF* mutant females exhibit intersex morphology**
(**A**) Images of control female (left), *dsxF* female (centre) and control male (right) anatomy. Side views of mosquito heads (top); images of mosquito heads (middle); images of genitals (bottom).
(**B**) Quantification of changes in flagellar lengths across control females, *dsxF* females and control males. Individual points represent lengths from individual mosquitoes. Solid lines represent means and standard deviations. Pairwise t-tests with BH corrections, *** $p<0.001$. See Tables S1 and S2 for measured values and statistical comparisons. Sample sizes: 45 control females; 24 *dsxF* females; 20 control males.

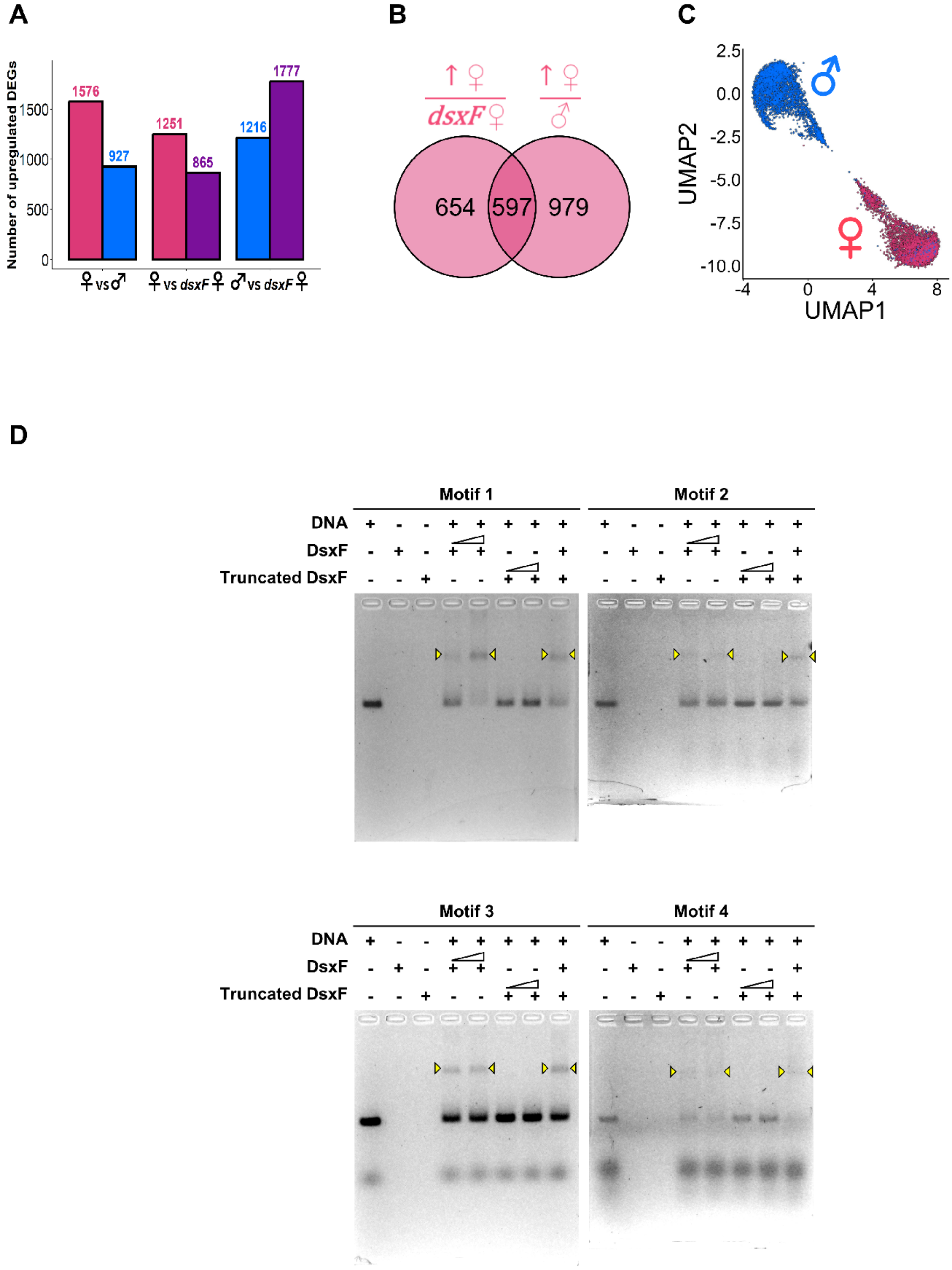

**Figure S2: Thorax RNA-sequencing analyses**
(**A**) Bar chart showing number of Differentially Expressed Genes (DEGs) for each comparison group. DEGs identified from DESeq2 analysis (padj<0.1, $log_2$FoldChange>0).
(**B**) Venn diagram of genes unregulated in the thorax of control females compared to that of *dsxF* females and control males.
(**C**) Sex-specific muscle cells identified based on the sex of the thorax tissue samples [1].
(**D**) Whole gel images of EMSA showing protein-DNA interactions between full-length DsxF protein or truncated DsxF protein without DM domain and the four 100 bp promoter fragments of *myo-fem* that each bears a centered Dsx-binding motif. Interactions represented as band shifts were observed only between full-length DsxF protein and target DNA sequences. See also Fig 1K.

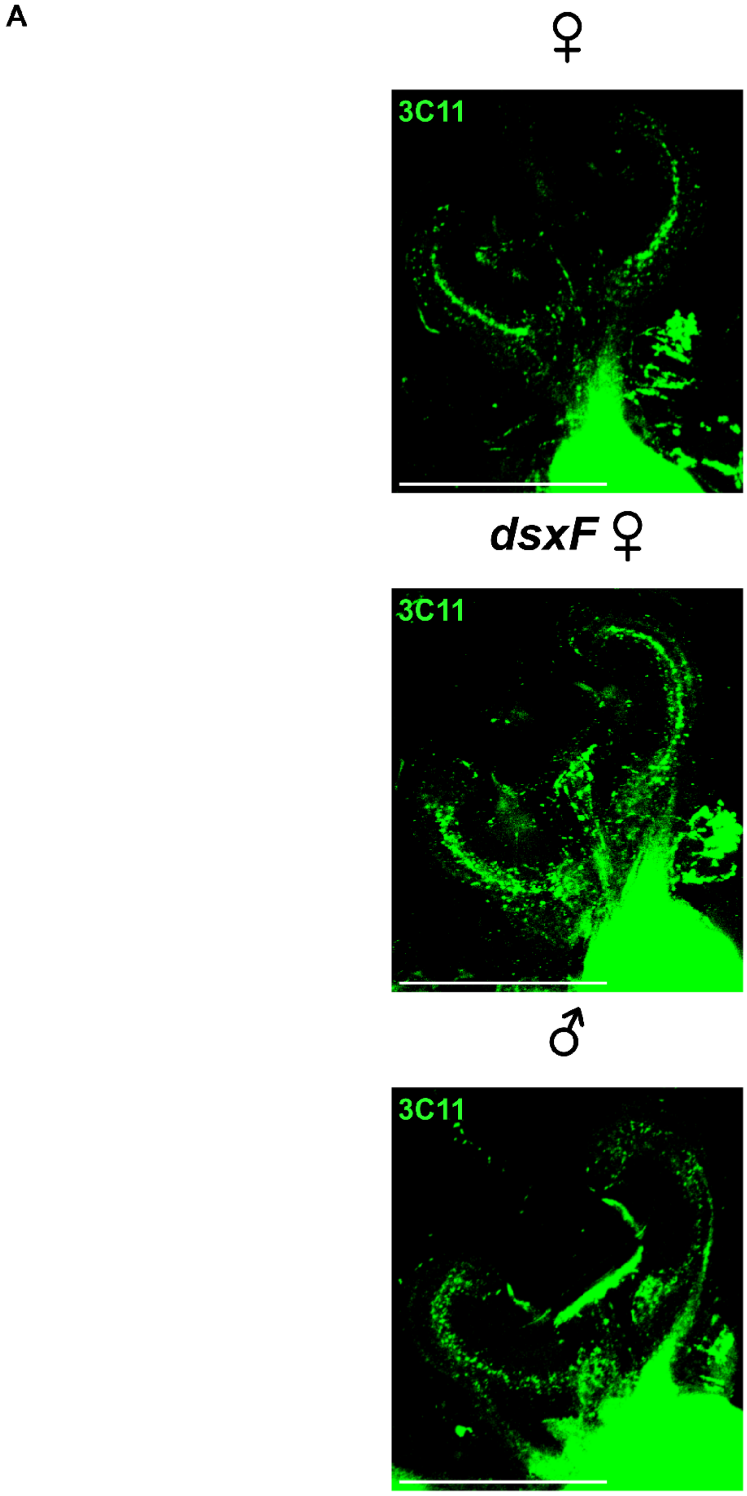

**Figure S3: Loss of DsxF alters mutant female JO neuroanatomy**
(**A**) anti-SYNORF1 (3C11) channel images of control female (top), *dsxF* female (middle) and control male (bottom) JO sections stained with anti-SYNORF1 (3C11, green). Scale bar, 100μm. See also Fig 2D.

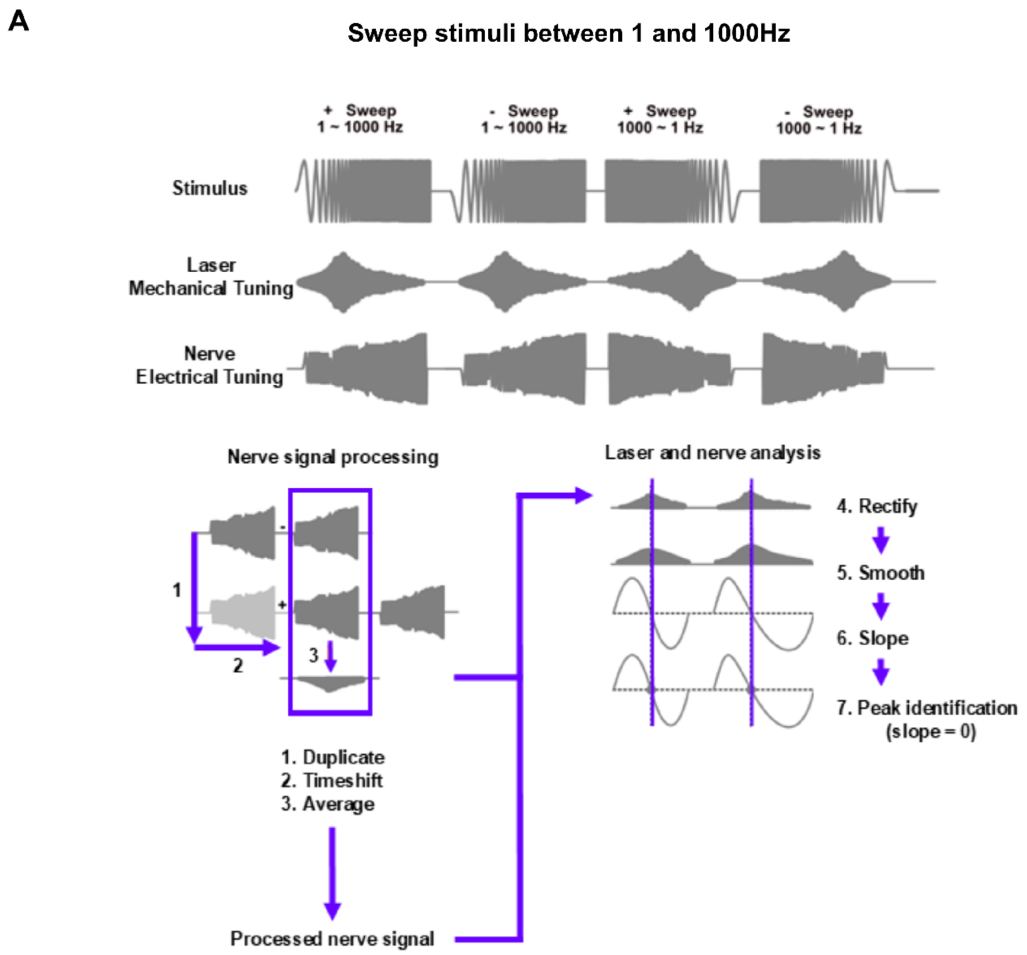

**Figure S4: Laser Doppler vibrometry coupled with electrophysiology sweep recordings analysis paradigm**

(**A**) Analysis paradigm for estimating mechanical and electrical tuning based on vibrometry/electrophysiology recordings. Sweep stimuli provided using electrostatic actuation (top), with flagellar displacement (middle) and antennal nerve compound action potential responses (bottom) simultaneously recorded. Nerve channel data was then duplicated, time-shifted and averaged to remove artefacts resulting from the use of electrostatic actuation. Processed nerve signals, and laser channel signals, were rectified and smoothed, before a slope function was applied. The point at which the slope was equal to zero (equivalent to the peak flagellar displacement/maximum antennal nerve response) was then identified.

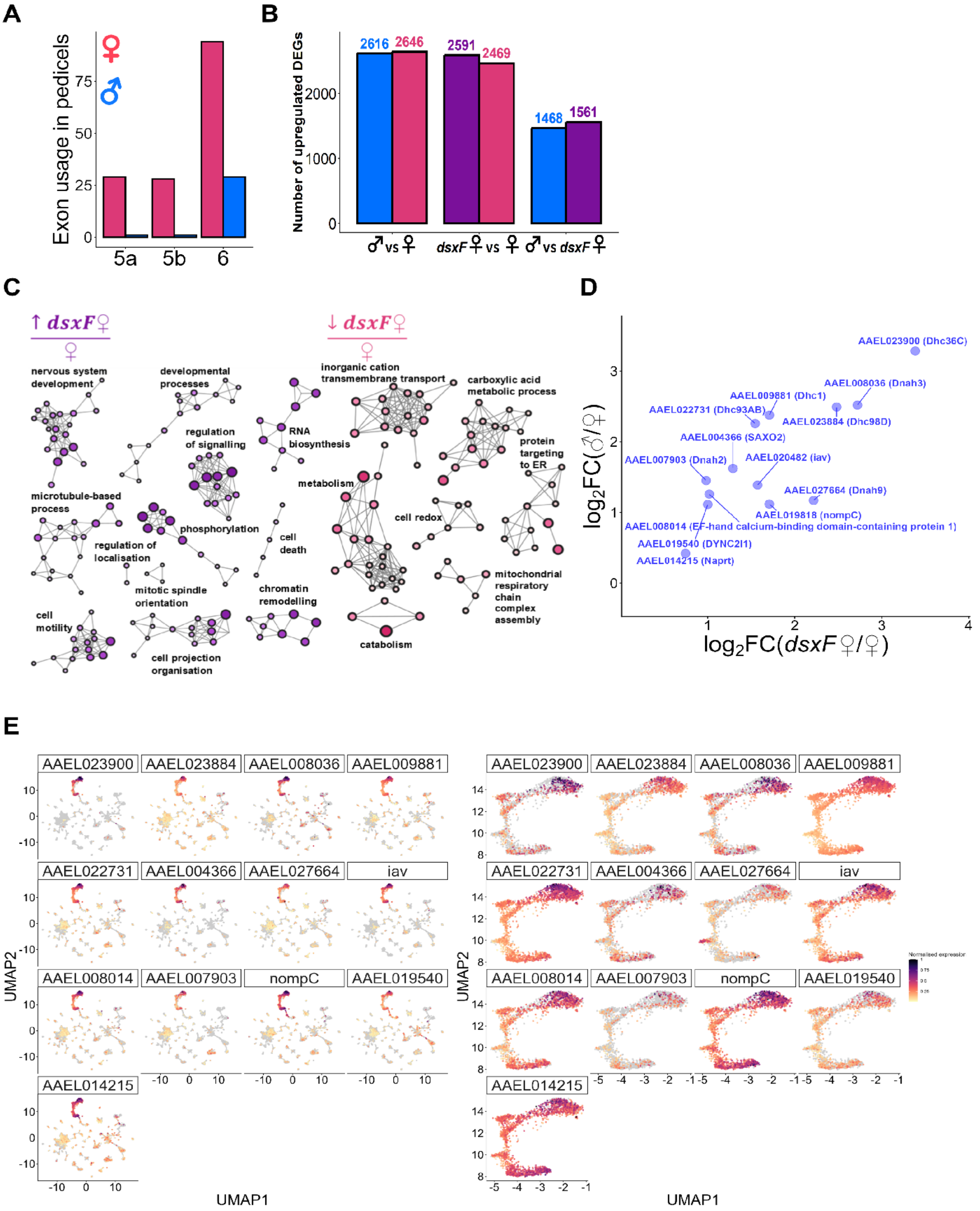

**Figure S5: Pedicel RNA-sequencing analyses**

(**A**) *dsx* splicing in control female and control male pedicels analysed based on published wild-type male and female pedicel bulk RNA-sequencing data [2]. Pink represents female exon usage, and blue represents male exon usage.

(**B**) Bar chart showing number of Differentially Expressed Genes (DEGs) for each comparison group. DEGs identified from DESeq2 analysis (padj<0.1, $\log_2$FoldChange>0).

(**C**) GO enrichment analysis for genes upregulated in *dsxF* females compared to control females (left) and downregulated in *dsxF* females compared to control females (right).

(**D**) $\text{Log}_2$FoldChange of intersection genes shown in Figs 4C and 4D in control male and *dsxF* female pedicels compared to the control females.

(**E**) Expression patterns of genes in (left) heads and (right) JO neurons. Data from previously published head snRNA-seq dataset [1]. Color intensity represents normalized UMI-count.

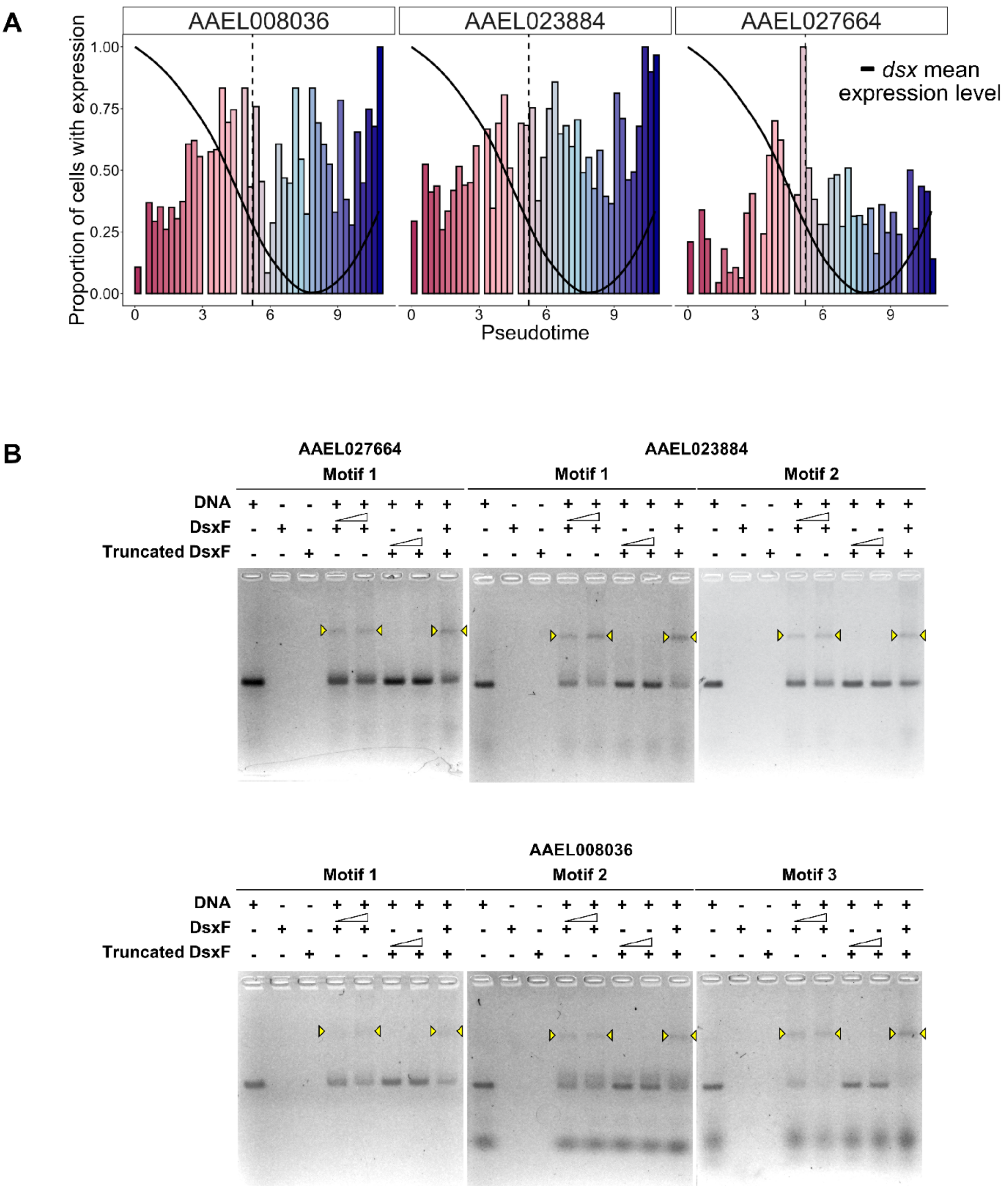

**Figure S6: DsxF directly inhibits expression of male-biased motile ciliary machinery genes in female pedicels**

(**A**) Changes in the proportion of nuclei expressing three different genes (AAEL008036, AAEL023884, AAEL027664) across *pseudotime* bins. Solid black line represents *dsx* mean expression level along the *pseudotime* trajectory. Color gradient represents change from female JO neurons to male JO neurons along the trajectory.

(**B**) Whole gel images of EMSA showing protein-DNA interactions between full-length DsxF protein or truncated DsxF protein without DM domain and the 100 bp promoter fragments from genes that each bears a centered Dsx-binding motif. Interactions represented as band shifts were observed only between full-length DsxF protein and target DNA sequences. See also Fig 4J.

### Supplemental Tables

**Table S1: Anatomical measurements for mosquito flagellae.** Values provided as means ± standard deviations. Numbers in brackets are sample sizes for each experiment.

| Group | Flagellum length (µm) |
|---|---|
| Control female | 1794.83 ± 63.53 (n = 45) |
| *dsxF* female | 1716.38 ± 90.41 (n = 24) |
| Control male | 1579.33 ± 32.17 (n = 20) |

**Table S2: Statistical values for flagellum length comparisons**. Pairwise t-tests with Benjamini Hochberg correction used for all comparisons.

| Comparison groups | Adjusted p-values | t-values | Degrees of freedom |
|---|---|---|---|
| Control male – Control female | $6.21 \times 10^{-26}$ | -18.12 | 61.79 |
| *dsxF* female – Control female | $5.76 \times 10^{-4}$ | -3.78 | 35.43 |
| *dsxF* female – Control male | $1.75 \times 10^{-7}$ | 6.919 | 29.69 |

**Table S3: JO width measurements.** Values provided as means ± standard deviations. Numbers in brackets refer to sample sizes for each experiment.

| Group | JO width (μm) |
|---|---|
| Control female | 135.41 ± 5.46 (n = 15) |
| *dsxF* female | 154.67 ± 7.09 (n = 14) |
| Control male | 173.13 ± 9.76 (n = 15) |

**Table S4: Statistical values for JO width comparisons.** Pairwise t-tests with Benjamini Hochberg correction used for all comparisons.

| Comparison groups | Adjusted p-values | t-values | Degrees of freedom |
|---|---|---|---|
| Control male – Control female | $2.35 \times 10^{-11}$ | 13.06 | 21.97 |
| *dsxF* female – Control female | $2.98 \times 10^{-8}$ | 8.15 | 24.41 |
| *dsxF* female – Control male | $3.86 \times 10^{-6}$ | -5.85 | 25.52 |

**Table S5: Stimulated mechanical and electrical tuning frequencies.** Data collected using vibrometry/electrophysiology paradigm utilising sweep stimuli. Values provided as means ± standard deviations. Numbers in brackets refer to sample sizes for each experiment.

| Group | Mechanical tuning - Sweep stimulation (Hz) | Electrical tuning - Sweep stimulation (Hz) |
|---|---|---|
| Control female | 245.25 ± 23.05 (n = 19) | 175.98 ± 22.35 (n = 19) |
| *dsxF* female | 349.13 ± 36.96 (n = 18) | 208.83 ± 21.95 (n = 18) |
| Control male | 405.42 ± 44.66 (n = 17) | 297.18 ± 45.0 (n = 17) |

**Table S6: Statistical values for mechanical tuning frequency during sweep stimulation comparisons.** Data collected using vibrometry/electrophysiology paradigm utilising sweep stimuli. Pairwise t-tests with Benjamini Hochberg correction used for all comparisons.

| Comparison groups | Adjusted p-values | t-values | Degrees of freedom |
|---|---|---|---|
| Control male – Control female | $6.69 \times 10^{-12}$ | 13.29 | 23.36 |
| *dsxF* female – Control female | $8.64 \times 10^{-11}$ | 10.20 | 28.22 |
| *dsxF* female – Control male | $3.16 \times 10^{-4}$ | -4.05 | 31.13 |

**Table S7: Statistical values for electrical tuning frequency during sweep stimulation comparisons**. Data collected using combined vibrometry and electrophysiology paradigm utilising sweep stimuli. Pairwise t-tests with Benjamini Hochberg correction used for all comparisons.

| Comparison groups | Adjusted p-values | t-values | Degrees of freedom |
|---|---|---|---|
| Control male – Control female | $2.23 \times 10^{-9}$ | 10.05 | 22.85 |
| *dsxF* female – Control female | $7.0 \times 10^{-5}$ | 4.51 | 34.95 |
| *dsxF* female – Control male | $2.96 \times 10^{-7}$ | -7.31 | 22.91 |

**Table S8: Unstimulated mechanical tuning frequencies.** Data collected using vibrometric measurements of unstimulated ear vibrations. Values provided as means ± standard deviations. Numbers in brackets refer to sample sizes for each experiment.

| Group | Mechanical tuning - Active unstimulated (Hz) | Mechanical tuning - Passive unstimulated (Hz) |
|---|---|---|
| Control female | 242.38 ± 8.70 (n = 15) | 234.37 ± 22.25 (n = 15) |
| *dsxF* female | 602.89 ± 58.96 (n =13) | 294.87 ± 27.76 (n = 13) |
| Control male | 664.01 ± 86.03 (n = 19) | 324.40 ± 27.67 (n = 19) |

**Table S9: Statistical values for mechanical tuning frequency in unstimulated, active and passive states comparisons.** Data collected using laser Doppler vibrometry. Pairwise t-tests used for all comparisons.

| Comparison groups | Adjusted p-values | t-values | Degrees of freedom |
|---|---|---|---|
| Control female active-passive | 0.17 | 1.44 | 14 |
| *dsxF* female active – passive | $4.16 \times 10^{-10}$ | 18.21 | 12 |
| Control male active - passive | $5.59 \times 10^{-12}$ | 15.77 | 18 |

**Table S10: Median values of maximum AUC ratio for Ringer and TeNT injection datasets (vibrometry).** All values provided as medians ± standard errors. Numbers in brackets refer to sample sizes for each experiment.

| Group | Maximum AUC ratio after Ringer injection | Maximum AUC ratio after TeNT injection |
|---|---|---|
| Control female | 1.40 ± 0.10 (n = 3) | 1.23 ± 0.12 (n = 3) |
| *dsxF* female | 1.48 ± 0.07 (n = 3) | 1.34 ± 0.24 (n = 3) |
| Control male | 1.66 ± 0.36 (n = 3) | 83500 ± 19812 (n = 3) |

**Table S11: Median values of maximum AUC ratio for Ringer and cAMP injection datasets (vibrometry).** All values provided as medians ± standard errors. Numbers in brackets refer to sample sizes for each experiment.

| Group | Maximum AUC ratio after Ringer injection | Maximum AUC ratio after cAMP injection |
|---|---|---|
| Control female | 1.24 ± 0.17 (n = 7) | 1.12 ± 0.19 (n = 10) |
| *dsxF* female | 1.41 ± 0.36 (n = 8) | 1.02 ± 0.50 (n = 7) |
| Control male | 1.72 ± 0.40 (n = 8) | 841.89 ± 3505.04 (n = 8) |

**Table S12: Statistical values for maximum AUC ratio comparisons for Ringer and cAMP injection datasets.** Data collected using laser Doppler vibrometry coupled with injections. Wilcoxon tests used for comparisons.

| Comparison groups | Adjusted p-values | W-values |
|---|---|---|
| Control female<br>Ringer - cAMP | 0.860 | 41 |
| *dsxF* female<br>Ringer - cAMP | 0.690 | 32 |
| Control male<br>Ringer - cAMP | $1.55 \times 10^{-4}$ | 0 |
